# Proteostatic tuning underpins the evolution of novel multicellular traits

**DOI:** 10.1101/2023.05.31.543183

**Authors:** Kristopher Montrose, Dung T. Lac, Anthony J. Burnetti, Kai Tong, G. Ozan Bozdag, Mikaela Hukkanen, William C. Ratcliff, Juha Saarikangas

## Abstract

The evolution of multicellularity paved the way for the origin of complex life on Earth, but little is known about the mechanistic basis of early multicellular evolution. Here, we examine the molecular basis of multicellular adaptation in the Multicellularity Long Term Evolution Experiment (MuLTEE). We demonstrate that cellular elongation, a key adaptation underpinning increased biophysical toughness and organismal size, is convergently driven by downregulation of the chaperone Hsp90. Mechanistically, Hsp90-mediated morphogenesis operates by destabilizing the cyclin-dependent kinase Cdc28, resulting in delayed mitosis and prolonged polarized growth. Reinstatement of Hsp90 or Cdc28 expression resulted in shortened cells that formed smaller groups with reduced multicellular fitness. Together, our results show how ancient protein folding systems can be tuned to drive rapid evolution at a new level of biological individuality by revealing novel developmental phenotypes.

**Teaser:** Downregulation of Hsp90 decouples cell cycle progression and growth to drive the evolution of macroscopic multicellularity.

## Introduction

The evolution of multicellular organisms from single-celled ancestors has independently occurred ∼50 times across the tree of life (*1–5*). Each of these events represents a major transition in individuality, but because they occurred in the deep past, relatively little information is available about the evolutionary dynamics and molecular mechanisms through which simple groups of cells evolve into multicellular organisms.

The transition to multicellularity may precipitate a period of rapid evolution, as cells adapt to novel organismal and ecological contexts (*6, 7*). Epigenetic mechanisms may play a crucial role in this process (*8*), as they are often capable of generating heritable phenotypic diversity at faster rates than mutation alone (*9–12*). In addition to mechanisms altering gene expression, many proteins exist in dynamic interconverting states of folding and assembly (*13–15*), which in some cases can produce heritable phenotypic variation that may serve as a basis for adaptive evolution (*16, 17*). However, given the ancient origins of extant multicellular clades, no work has directly examined the role of epigenetic inheritance in the evolution of multicellularity. Our experiment aims to circumvent this constraint through long-term directed evolution, providing insights into the potential role of non-genetic mechanisms during the early stages of multicellular transition.

Using long-term experimental evolution to select for larger size over thousands of generations, we recently showed that multicellular ‘snowflake yeast’ can evolve to form multicellular groups that are over 20,000 times larger and 10,000 times more mechanically tough than their ancestors (*18*). Cellular elongation played a central role in the evolution of these novel multicellular traits, allowing branches of cells to entangle with one another and thereby become orders of magnitude more mechanically tough (*19*). Here we set out to investigate the underlying molecular mechanisms behind cellular elongation and macroscopic multicellularity. We found that downregulation of the chaperone protein Hsp90, a key modulator of genotype-phenotype relationships, was a convergent adaptation underpinning the evolution of larger, more mechanically tough groups. Mechanistically, we found that reduced Hsp90 expression leads to modulation of cell morphogenesis by destabilizing the cyclin-dependent protein kinase Cdc28, resulting in delayed cell cycle progression and consequent cellular elongation. Collectively, these data unravel that altered chaperoning of cellular proteome can facilitate major evolutionary transitions by generating novel cell-level phenotypic traits that promote multicellular evolution.

## Results

### Hsp90 is downregulated during evolution of macroscopic multicellularity

Our ongoing Multicellularity Long Term Evolution Experiment (MuLTEE) allows us to examine multicellular evolution in a nascent lineage of clonally-developing organisms. This experiment was initiated in *S. cerevisiae* strains lacking the *ACE2* open reading frame to generate small ‘snowflake’ cell clusters (*20, 21*) referred hereafter as ‘Ancestors’. During the initial 600 rounds of size-based selection (∼3,000 generations) for aerobic, mixotrophic, and anaerobic snowflake yeast populations, only the anaerobic lines evolved macroscopic size (**Fig. 1A**) (18). This multicellular adaptation was the result of a morphological transformation from oval to rod-shaped cells, leading to a significant increase in cellular aspect ratio (ratio of length to width) (**Fig. 1B, 1C**). Elongated cells form long branches which mutually entangle, resulting in the evolution of far tougher multicellular groups (i.e., from weaker than gelatin to as strong as wood), allowing individual snowflake yeast clusters to grow to macroscopic sizes (**Fig. 1D**) (18).

**Figure 1:**
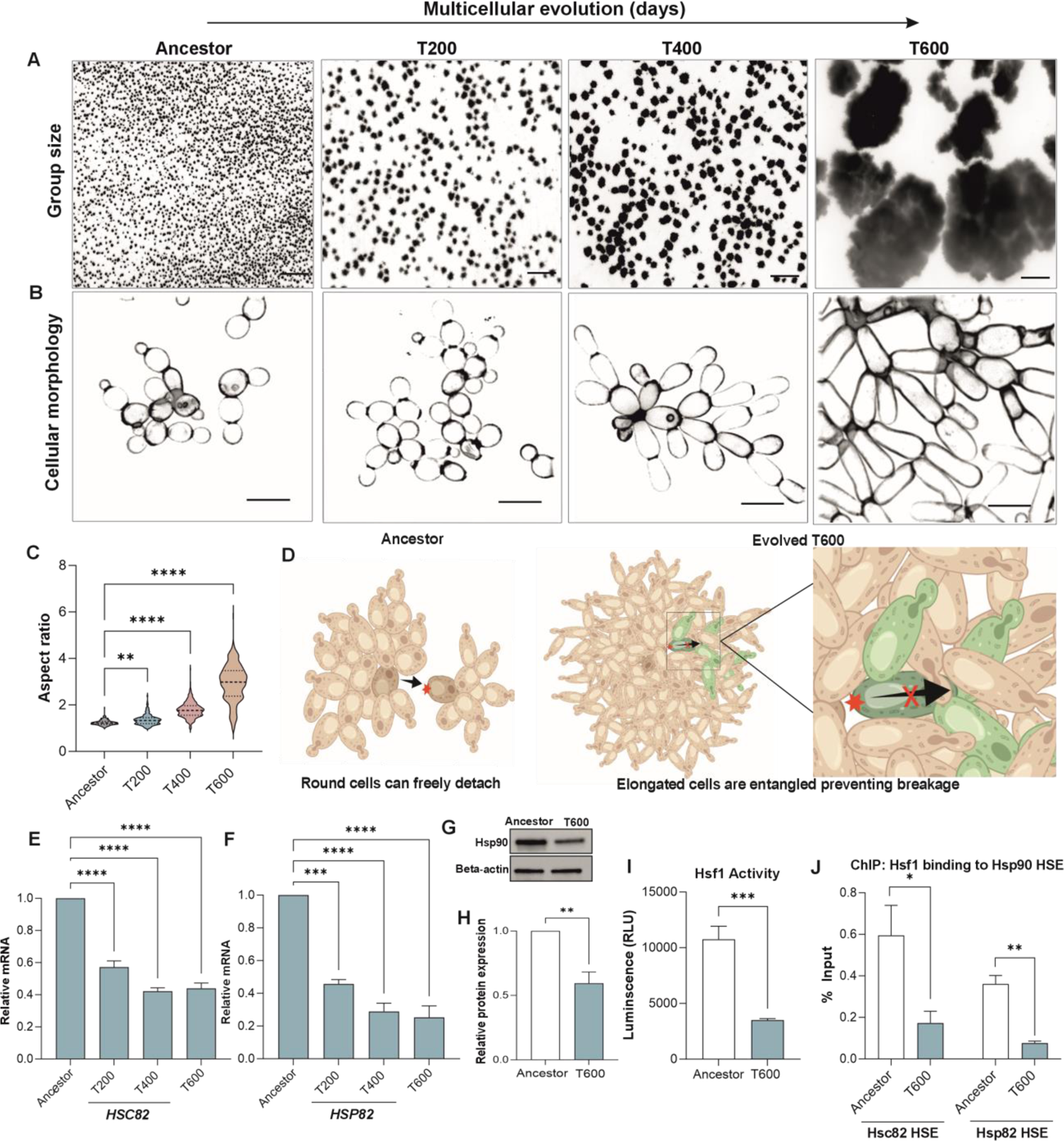
Hsp90 is downregulated during the evolution of macroscopic multicellularity in snowflake yeast. (A) Representative images of Snowflake yeast clusters at 200-day time points during the experimental evolution experiment selecting for larger group size (MuLTEE). Scale bar = 500 µm. (B) Representative images of Snowflake yeast cell morphology at 200-day time points during the MuLTEE. Scale bar = 10 µm. (C) Quantification of the cellular aspect ratio of snowflake yeast at 200 day time points (*n*=300 cells, *F*_3, 1196_ = 220.5, *p* < 0.0001, one way ANOVA, *Tukey’s* post hoc test Ancestor vs T200 *p* = 0.004, Ancestor vs T400 *p* < 0.0001, Ancestor vs T600 *p* < 0.0001 one way ANOVA). (D) Graphic depicting the effect of cellular elongation on increasing entanglement, leading to clusters that are robust to fracture (*19, 21*). (E) Quantification of *HSC82* expression levels by RT-qPCR in T200, T400 and T600 compared to Ancestor (*n*=3, *F*_3, 8_ = 96.65, *p* < 0.0001, one way ANOVA, *Tukey’s* post hoc test Ancestor vs T200 *p* < 0.0001, Ancestor vs T400 *p* < 0.0001, Ancestor vs T600 *p* < 0.0001, one way ANOVA). (F) Quantification of *HSP82* expression levels by RT-qPCR in T200, T400 and T600 compared to Ancestor (*n*=3, *F*_3, 8_ = 43.16, *p* < 0.0001, *Tukey’s* post hoc test Ancestor vs T200 *p* < 0.0001, Ancestor vs T400 *p* < 0.0001, Ancestor vs T600 *p* < 0.0001, one way ANOVA). (G) Representative immunoblot of Hsp90 expressed by Ancestor and T600 detected using an antibody against Hsp90. Antibody against Beta-actin was used as a loading control. (H) Quantification of relative band intensity of T600 Hsp90 in comparison to Ancestor Hsp90. Bands were normalized to Beta actin loading control (*n*=3, *t* = 4.6, *p* = 0.0098, two-sample t-test). (I) Hsf1 activity in Ancestor and T600 measured as luminescence. Measurement readings of substrate alone were subtracted as background (*n*=4, *t* = 6.1, *p* < 0.0009, two-sample t-test). (J) Hsf1 ChIP-QPCR analysis of Hsc82 and Hsp82 HSE regions in Ancestor and T600. Background was determined using controls with no antibody. Data was normalized against input signal (*n*=4, Hsc82 *t* = 2.7, *p =* 0.026, Hsp82 *t* = 5.8, *p* = 0.002, two-sample *t*-test). All values represent mean ± SEM.

To identify the molecular changes that underlie the morphological transformation from oval-shaped Ancestor cells to the rod-shaped T600 cells, we examined the transcriptomes of anaerobic line 5 (PA5), which grew the largest clusters after 600 days of settling selection. Our results identified ∼540 downregulated genes and ∼460 upregulated genes (*p* < *0.05*, log2 fold change cut-off 0.5) in the T600 cells compared to Ancestors. Examining the top 50 gene ontology (GO) terms revealed that the major differentially expressed genes were involved in metabolism, cell cycle, and translation (**Fig. S1A**). Interestingly, we found that a number of chaperone proteins were downregulated in T600 cells including both isoforms of yeast Hsp90, Hsc82, and Hsp82 (**Fig. S1B**). Hsp90 is particularly interesting as it is involved in the final folding steps of specialized client proteins that include transcription factors and kinases, and thereby is controlling their activity post-translationally. Thus, it can have a significant impact on the genotype-phenotype relationship by altering the activity of key developmental pathways (*22, 23*).

We confirmed the RNAseq results using quantitative PCR, which demonstrated that *HSC82* mRNA was reduced by ∼2 fold and *HSP82* mRNA by ∼3 fold in T600 cells as compared to the Ancestor (**Fig. 1E, 1F**). Further analysis revealed that decline in Hsp90 mRNA expression can be already observed starting from day 200 of evolution. Consistent with declining mRNA expression, Western blot analysis using an antibody that recognizes both Hsp90 isoforms revealed that Hsp90 declined by ∼40% in T600 cells, relative to the Ancestor (**Fig. 1G, 1H**).

To determine the cause of Hsp90 decline, we investigated the activity of transcription factor Hsf1, which is responsible for the expression of yeast Hsp90 (*24*). Hsf1 binds to heat shock elements (HSEs) found in the promoter region of its target genes, such as Hsp90, to activate transcription. In yeast, Hsf1 is active at basal levels promoting transcription of constitutively expressed Hsc82 and it can be further induced by stress, such as heat, to promote the expression of stress-induced Hsp82. We measured the activity of Hsf1 by quantifying the expression of a genomically-integrated luciferase reporter gene that is under the control of HSE promoter regions (*25*). Using luminescence output as a measure of Hsf1 activity, we found that there was less Hsf1 activity occurring in T600 cells compared to Ancestor cells at both the basal level (**Fig. 1I**) and after heat shock (**Fig. S1C**). Our whole genome sequencing detected no mutations in *HSF1*, the HSE regions of *HSP82* or *HSC82*, or in other regulators of the heat shock response. Additionally, there were no differences in the expression of *HSF1* between the Ancestor and T600 cells (**Fig. S1D**). To test if the altered Hsf1 activity is mediated by reduced target gene binding, we used a chromatin-immunoprecipitation (ChIP) assay to analyze the binding of Hsf1 to Hsp90 HSE promoter regions. Compared to the Ancestor, we found a ∼70% reduction in Hsf1 occupancy at Hsp90 HSE regions in T600 (**Fig. 1J**), explaining the loss of activity and reduced Hsp90 expression. Together, these results indicate that Hsp90 downregulation is mediated by its reduced transcription by Hsf1.

### Downregulation of Hsp90 plays a critical role in snowflake yeast evolution

We next sought to determine whether the reduced Hsp90 expression is a multicellular adaptation, giving rise to novel, beneficial multicellular traits in evolved snowflake yeast. To investigate this, we restored the expression level of Hsp90 back to the ancestral level by integrating *TEF1pr-HSC82* into a safe harbor locus in the snowflake genome (T600-Hsc82OE). This led to a ∼50% increase in *HSC82* mRNA expression (**Fig. S2A**). Interestingly, restoring *HCS82* levels led to cells reverting to shorter and rounder morphology (**Fig. 2A**), with a significant reduction in mean aspect ratio from 2.93 in wildtype T600 cells to 2.18 in T600-Hsc82OE (**Fig. 2B**). To further validate that a reduction in Hsp90 expression affects cell shape, we subjected Ancestor cells to a short-term treatment with radicicol, a chemical inhibitor of Hsp90. This had a modest but significant effect on the Ancestor, increasing the cellular aspect ratio by ∼10% after just 6 h of growth (**Fig. 2C**).

**Figure 2:**
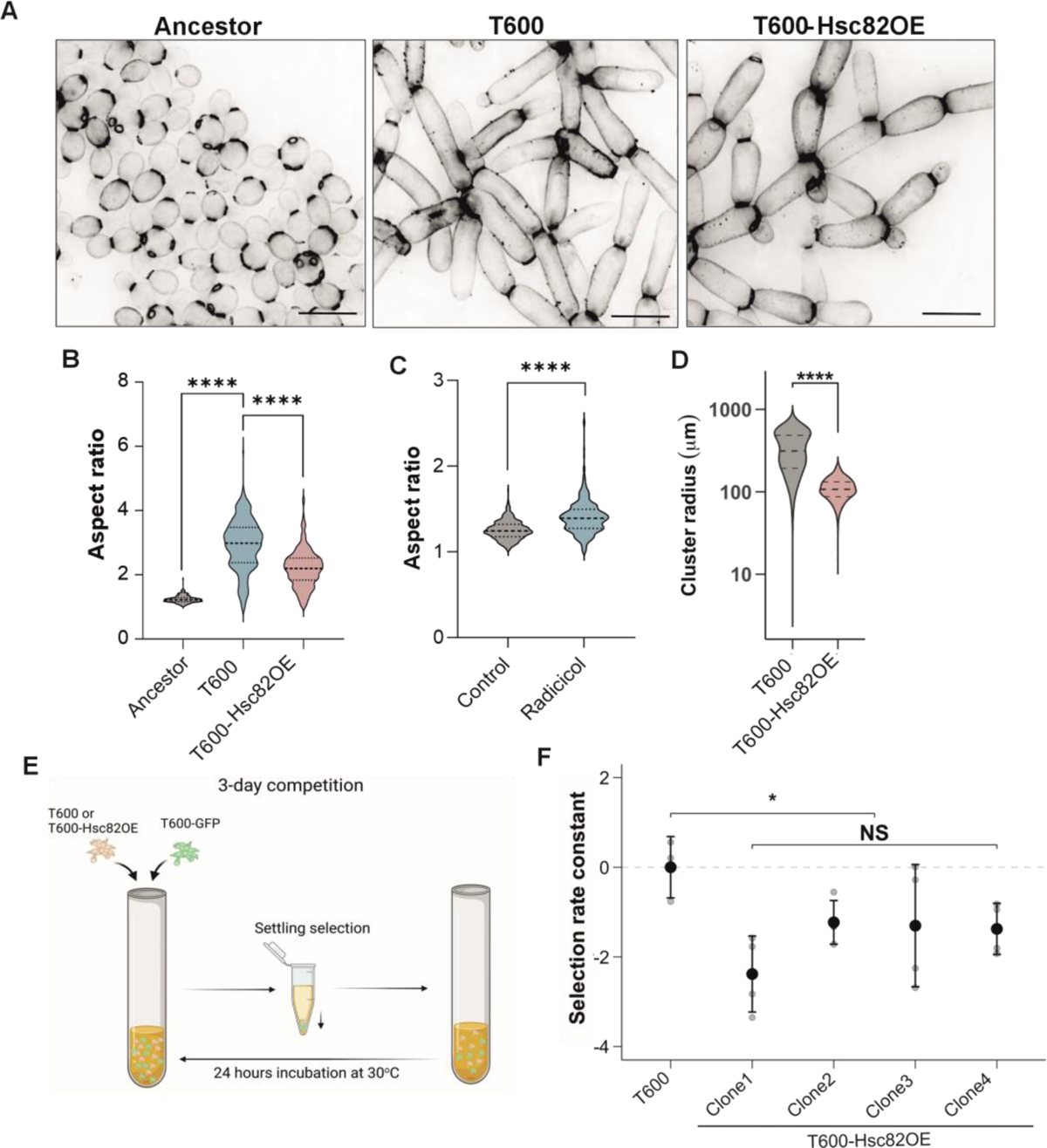
Overexpression of *HSC82* decreases aspect ratio and cluster size of T600 cells. (A) Representative images of Ancestor, T600 and T600-Hsc82OE cells, highlighting differences in cellular morphology. Scale bar = 10 µm. (B) Quantification of the cellular aspect ratio of Ancestor, T600 and four T600-Hsc82OE clones combined (*n*=300 cells, T600-Hsc82OE. *F*_2,897_ = 665.9, *p* < 0.0001, one way ANOVA, *Tukey’s* post hoc test Ancestor vs T600 *p* < 0.0001, T600 vs T600-Hsc82OE *p* < 0.0001). (C) Quantification of the cellular aspect ratio of Ancestor and radicicol-treated Ancestor (*n*=300 cells, *t =* 16.1, *p* < 0.0001, two-sample t-test). (D) Cluster size as a measure of cluster radius (µM) for T600 and T600-Hsc82OE (T600 *n*=1033, T600-Hsc82OE *n*=3489 from 4 clones), (*F*_1,6229_ = 3024, *p* < 0.0001, one way ANOVA, *Tukey’s* post hoc test T600 vs Hsc82OE *p* < 0.0001). (E) Diagram summarizing the competition assay method in (F). All strains were competed against a T600-GFP. (F) Fitness over three rounds of growth and selection, represented as a selection rate constant, for the T600 isolate and four independently-generated clones of T600-Hsc82OE (*n*=4 per competition). To account for the cost of GFP expression, all selection rate constants were normalized by the mean of the T600 vs. T600-GFP competition. (*n*=3 per competition) (*F*_1, 14_ = 8.4, *p* = 0.015, nested ANOVA). All values represent mean ± SEM.

We then tested whether the increased aspect ratio created by reduced Hsp90 expression is adaptive under our selective regime (24 h of growth in shaking incubation, followed by settling selection favoring larger groups). Our previous work has established that cellular elongation is a key trait underpinning the origin of increased multicellular size and toughness, convergently evolving in all five replicate populations evolving macroscopic size (*18, 26*). As expected, overexpressing Hsp90 dramatically reduced cluster size in four independently-generated PA5 T600-Hsc82OE clones, with a mean 3.7 fold reduction in size (**Fig. 2D, S2B**). We also examined the impact of increased Hsp90 expression on fitness via competition experiments (**Fig. 2E**). Restoring Hsp90 to ancestral levels was costly, resulting in a mean selection rate constant of −1.57 for the 3 days of competition (**Fig. 2F**). Together, these results establish that novel regulation of Hsp90 is an adaptive multicellular trait, increasing cellular aspect ratio and thus multicellular size and fitness.

### Convergent evolution of reduced Hsp90 expression

Macroscopic cluster size evolved convergently across the five replicate populations of our anaerobic yeast (PA1-5) through cellular elongation and entanglement (*19*). To determine if this trait is driven by reduced Hsp90 expression in all replicate populations, we examined *HSC82* expression as they were evolving large size (transfers 0-400). All five lines showed significant declines in *HSC82* at T200 and T400, relative to the Ancestor (**Fig. 3A**). Moreover, changes in the Hsp90 expression level, cellular aspect ratio (**Fig. 3B**), and cluster size (**Fig S3A**) correlated. Replicate lines 2, 4, and 5 show the steepest drop in Hsp90 levels as multicellular evolution progressed. Correspondingly, these populations led to the largest and most rapid change in cellular aspect ratio and cluster size (**Fig. 3B, S3A**).

**Figure 3:**
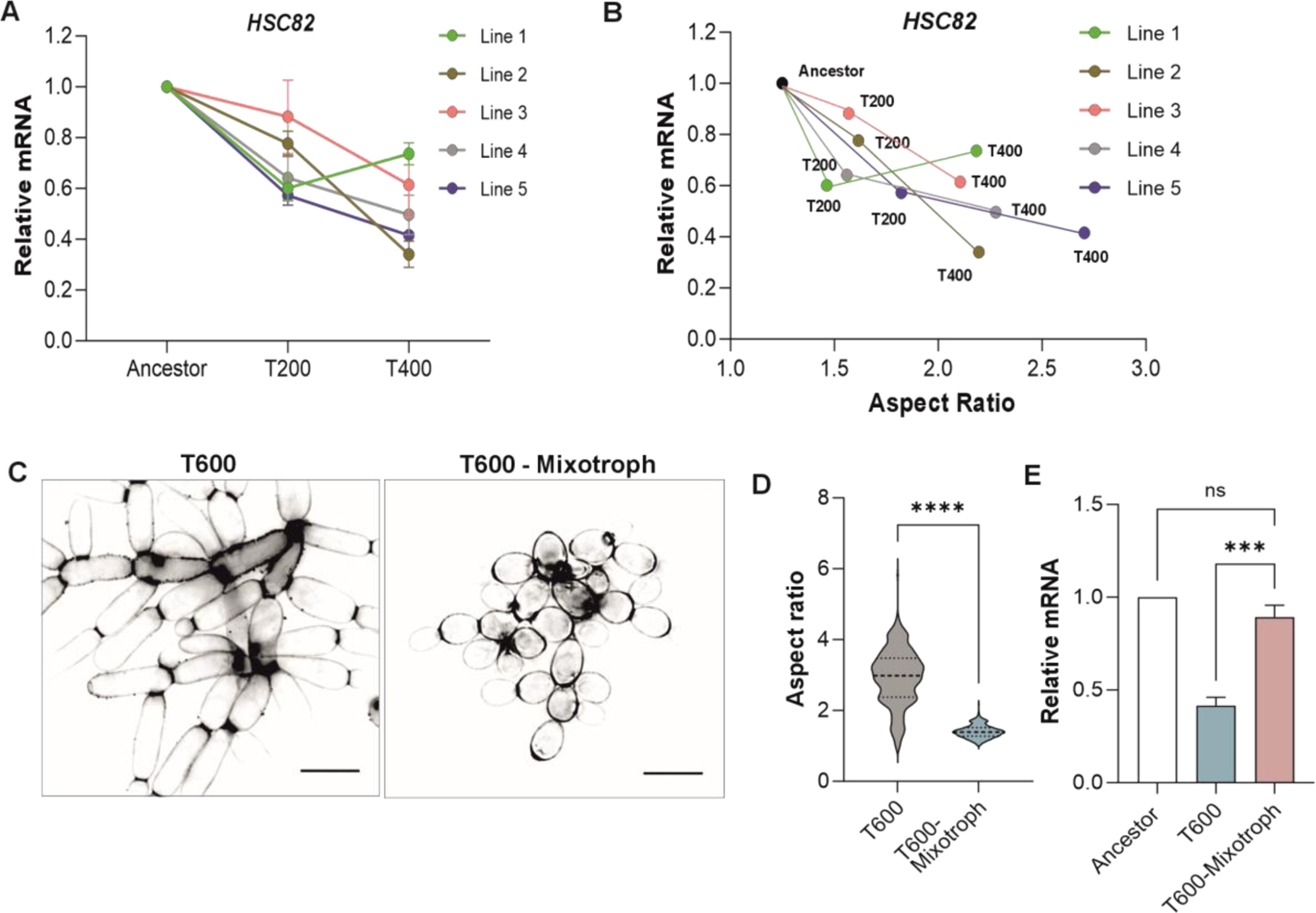
Expression of *HSC82* affects the timing and extent of macroscopic development. (A) Quantification of *HSC82* expression level by RT-QPCR in T200 and T400 compared to Ancestor for the five independent lines of aerobic snowflake yeast (*n*=4, *F_8_*_,20_ = 4.485, *p* = 0.003, two way ANOVA, *Tukey’s* post hoc test Ancestor vs T200 *p* < 0.0001, Ancestor vs T400 *p* < 0.0001, T200 vs T400 *p* = 0.025). (B) *HSC82* expression against aspect ratio for T200 and T400 of each of the five lines of aerobic snowflake yeast, (*r* = 0.74, *p* = 0.009, *y = −*0.33*x+*1.26, Linear regression). (C) Representative images of T600 and T600-Mixotroph to highlight differences in cellular morphology. Scale bar = 10 µm. (D) Quantification of the cellular aspect ratio of T600 and T600-Mixotroph (*n*=300 cells, *t =* 32.58, *p* < 0.0001, two-sample t-test). (E) Quantification of *HSC82* expression levels by RT-QPCR in T600 and T600-Mixotroph compared to Ancestor (*n*=3, *F*_2, 6_ = 48.33, *p* = 0.002, one way ANOVA, *Tukey’s* post hoc test Ancestor vs T600-mixotroph *p* = 0.28, T600-mixotroph vs T600 *p* < 0.0007). All values represent mean ± SEM.

Finally, we wanted to determine whether reduced Hsp90 expression is simply a byproduct of ∼3,000 generations of evolution, and not directly related to the evolution of larger group size. To this end, we measured Hsp90 expression levels in mixotrophic snowflake yeast, a separate treatment of the MuLTEE that was begun with snowflake yeast capable of aerobic metabolism. These yeast underwent the same 600 days of settling selection, but did not evolve macroscopic size or elongated cellular morphology (**Fig. 3C, 3D**). Prior work on this system has shown that, under our laboratory conditions, the ability to use oxygen for growth inhibits the evolution of large size. This is because oxygen diffuses poorly through the cluster and is mainly used by surface cells, incurring a size-dependent growth cost that is absent in anaerobic populations (*18, 19*). In contrast to the macroscopic strains which evolved under anaerobic metabolism, the mixotrophic T600 strain expressed *HSC82* at the same level as the anaerobic Ancestor and displayed approximately 50% higher expression than the anaerobic PA5 T600 strain (**Fig. 3E**). Collectively, these results provide evidence that downregulation of Hsp90 has occurred convergently to facilitate the evolution of macroscopic multicellularity.

### Loss of Hsp90 client Cdc28 leads to adaptive morphogenesis of evolved snowflake yeast

Cdc28 is the catalytic subunit of the yeast cyclin dependent kinase that acts as a master regulator of mitotic cell cycle through multiple combinatorial effects (*27*). Cdc28 is also a folding client of Hsp90 (*28, 29*). Inhibiting Hsp90 can alter Cdc28 protein expression, which in turn is linked to morphological changes (*28–32*). We examined whether Cdc28 is the downstream target of Hsp90 responsible for driving the elongated cell morphology of T600 cells. Our RNAseq showed no significant difference in *CDC28* mRNA expression between Ancestor and T600 cells, which we confirmed by qPCR (**Fig. 4A**). However, Cdc28 protein expression was ∼25% lower in T600 cells (**Fig. 4B-4C**). We then investigated whether the reduced stability of Cdc28 protein was caused by reduced Hsp90 expression by examining whether overexpressing Hsc82 increased Cdc28 abundance. We found that Hsc82 over-expression in T600 cells restored ∼80% of the reduced Cdc28 protein expression that evolved over 600 transfers (**Fig. 4C**), demonstrating that it is a target of Hsp90 that becomes destabilized in the T600 cells due to reduced Hsp90 expression.

**Figure 4:**
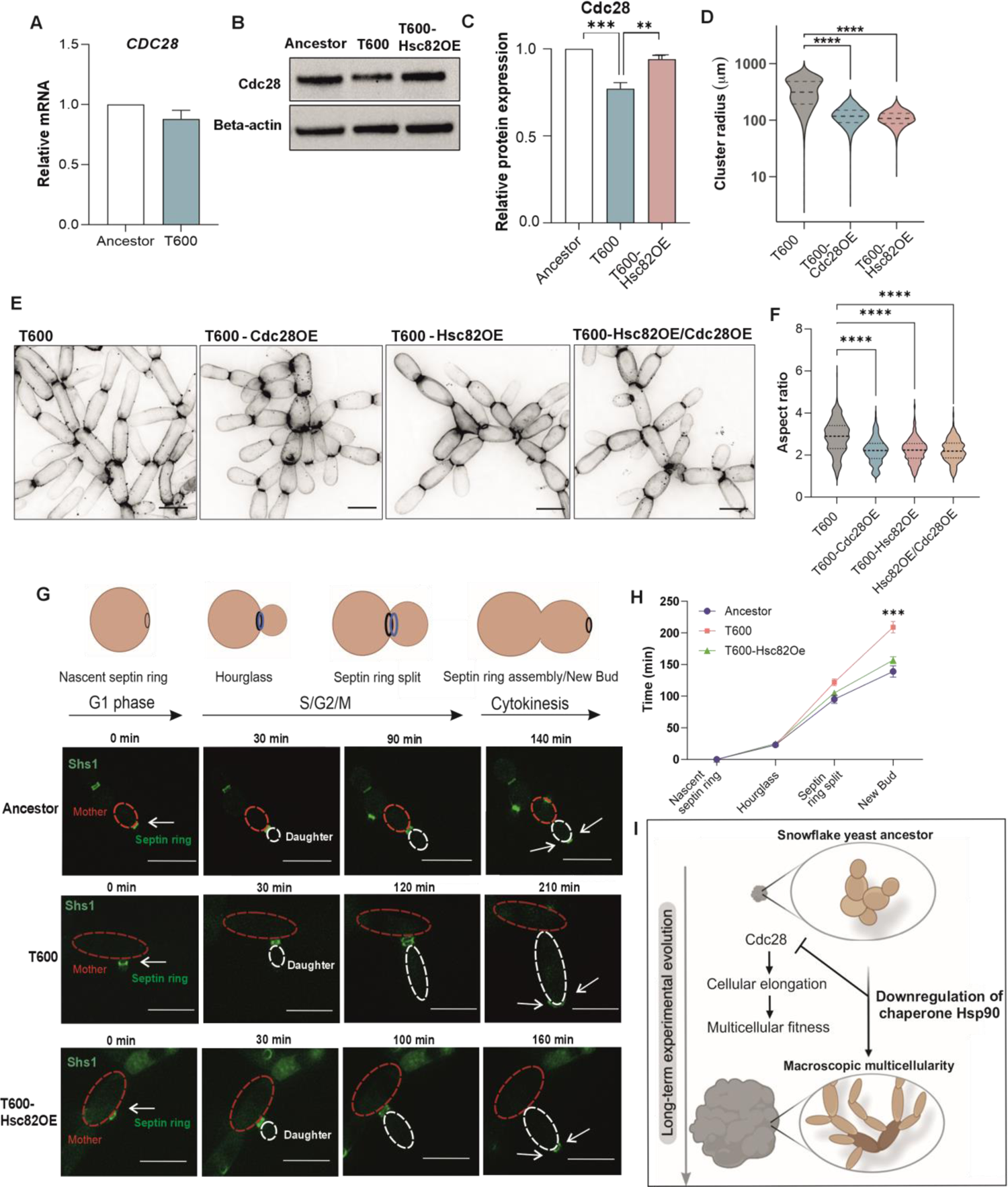
Elongation of T600 cells is driven by Hsc82 mediated destabilization of Cdc28. (A) Quantification of *CDC28* expression level by RT-qPCR of T600 compared to Ancestor (*n*=4, *t =* 1.67, *p* = 0.16, two-sample t-test). (B) Representative immunoblot of Cdc28 expressed by Ancestor, T600 and T600-Hsc82OE, detected with anti-Cdc28 antibody. Antibody against Beta-actin was used as a loading control. (C) Quantification of relative band intensity of T600 Cdc28 and T600-Hsc82OE Cdc28 in comparison to Ancestor Cdc28. Bands were normalized to loading control (*n*=4, *F*_2,9_ = 22.35, *p* = 0.0003, one way ANOVA, *Tukey’s* post hoc test Ancestor vs T600 *p* = 0.0003, T600 vs T600-Hsc82OE *p* = 0.0025). (D) Cluster size as a measure of cluster radius (µM) for T600, T600-Cdc28OE and T600-Hsc82OE (T600 *n*=2742, T600-Cdc28OE *n*=4458 from 4 clones, T600-Hsc82OE *n*=3489 from 4 clones, *F*_1,7195_ = 3477, *p* < 0.0001, one way ANOVA, *Tukey’s* post hoc test T600 vs T600-Cdc28OE *p* < 0.0001). (E) Representative images of T600, T600-Cdc28OE, T600-Hsc82OE and T600-Hsc82OE/Cdc28OE cells to highlight cellular morphology. Scale bar = 10 µm. (F) Quantification of the cellular aspect ratio of T600, T600-Cdc28OE, T600-Hsc82OE and T600-Hsc82OE/Cdc28OE (*n*=400 cells, T600-Cdc28OE and T600-Hsc82OE from 4 combined clones, T600-Hsc82OE/Cdc28OE from 3 combined clones. *F_3_*_,1596_ = 25.25, *p* < 0.0001, one way ANOVA, *Tukey’s* post hoc test T600 vs T600-Cdc28OE *p* < 0.0001, T600 vs T600-Hsc82OE *p* < 0.0001, T600 vs T600-Hsc82OE/Cdc28OE *p* < 0.0001). (G) Representative images of mNeonGreen-tagged Shs1 expressed by Ancestor, T600 and T600-Hsc82OE. Mother cell highlighted with red dashed line and daughter cell with white dashed line. Scale bar = 10 µm. Graphic summarizes relationship between septin ring stage and cell cycle stage. (H) Graphical representation of time lapse images for Ancestor, T600 and T600-Hsc82OE (*n*=10, *F*_6,72_ = 14.52, *p* < 0.0001, two way ANOVA, *Tukey’s* post hoc test, New Bud Ancestor vs New Bud T600 *p* = 0.0002, New Bud T600 vs New Bud T600-Hsc82OE *p* = 0.0008, New Bud Ancestor vs New Bud T600-Hsc82OE *p =* 0.18). (I) Graphical model for how downregulation of Hsp90 acts on Cdc28 to delay the cell cycle leading to the cellular elongation required for macroscopic evolution during the MuLTEE. All values represent mean ± SEM.

Hsp90 has hundreds of client proteins (*33–35*). To test whether the novel regulation of Cdc28 is responsible for the cellular elongation phenotype downstream of Hsp90, we inserted a single copy of *CDC28* under its own promoter in a safe harbor locus in the T600 strain (T600-Cdc28OE), which restored the expression of Cdc28 close to that of the Ancestor strain (average relative expression of T600-Cdc28OE was 1.25 ± 0.14 times that of the Ancestor strain from four clones) (**Fig. S4A-B**). Notably, this minor elevation in Cdc28 levels resulted in a significantly smaller cluster size, resembling that of T600-Hsc82OE (**Fig. 4D, S4C**). We then examined whether this effect was due to changes in cellular morphology by measuring the cellular aspect ratio. This showed that the restoration of Cdc28 expression resulted in cells with a smaller, rounder morphology, phenocopying the cellular aspect ratio of *HSC82* overexpressing T600 cells (**Fig. 4E-F**). Finally, to determine whether Hsc82 and Cdc28 exert their effects on cellular elongation through the same pathway, we conducted an epistasis analysis by simultaneously restoring the expression of both genes. Importantly, combined over-expression did not show additive effects on cellular aspect ratio compared to the single overexpression of either gene, demonstrating that Cdc28 is a key downstream factor in Hsp90-mediated control of cellular elongation in T600 cells (**Fig. 4F**).

Cell cycle and cell growth are loosely coupled (*36*). We postulated that the Hsp90-mediated reduction of Cdc28 may lead to excessive polarized growth due to delay in mitotic progression, as shown for some *CDC28* mutants (*30, 37*). To investigate this, we analyzed cell cycle dynamics by using a mNEONgreen-tagged copy of the septin ring subunit Shs1 as a proxy for cell cycle progression (*38*). T600 cells showed a delayed mitosis with slower progression of septin-ring splitting and the late mitotic events relative to Ancestor cells (**Fig. 4G, 4H**). To investigate if this was due to downregulated Hsp90, we overexpressed *HSC82* in T600 cells. This accelerated the cell cycle, and T600 cells were now progressing through mitosis at nearly the speed of Ancestor cells, providing evidence that Hsp90-mediated adaptive cellular morphogenesis in T600 is coupled to novel control of the cell cycle. These results provide evidence that adaptive multicellular morphogenesis established by the Hsp90-Cdc28-axis acts through delaying the cell cycle kinetics, enabling cells to undergo prolonged polarized growth that leads to their elongation (**Fig. 4I**).

## Discussion

In this work, we investigated the molecular basis of multicellular adaptation during the Multicellularity Long Term Evolution Experiment (MuLTEE). We found that downregulation of the chaperone protein Hsp90 was a convergent and adaptive trait that drove cellular elongation by modulating the stability and activity of the central cell cycle kinase Cdc28. This delayed progression through the cell cycle, resulting in prolonged polarized growth and the formation of elongated cells, which generate biomechanically-tough multicellular groups with higher Darwinian fitness. Our results thus reveal how manipulation of ancient systems that guide protein folding can facilitate major evolutionary transitions by generating novel developmental phenotypes through proteostatic tuning.

Previous research has established that Hsp90 can influence phenotypic variation and evolvability in a broad range of multicellular organisms by buffering or revealing cryptic genetic variation (*39–43*). Most prior work has examined how environmental stress acts as a catalyst for changes in Hsp90 function, subsequently leading to the appearance of Hsp90-dependent phenotypic alterations. In contrast, we demonstrate that Hsp90 can be under long-term selection for its role in regulating gene expression, generating novel, adaptive traits by modifying the activity of key client proteins. Our results are in line with a recent study showing that the introduction of a heterologous copy of Hsp90 from evolutionarily divergent *Y. lipolytica* to *S.cerevisiae* broadened the phenotypic space for natural selection (*44*). Together, these studies establish that Hsp90 function can be evolutionarily tuned to facilitate rapid adaptation.

At first glance, the loss of Hsp90 activity appears to contradict the broader macroevolutionary trend, in which increasingly complex organisms evolve increasingly complex proteomes (*45*). However, this largely occurs through an increase in the number of co-chaperones, rather than core chaperones, some of which have actually been lost prior to major evolutionary transitions. Comparative analysis of genes lost in animals revealed that Hsp100/ClpB chaperones are present in the closest-living relatives of animals, but are lost at the base of metazoa (*45, 46*). Such loss of core chaperones prior to the rapid diversification of animals might indicate an adaptive role for altered proteostatic tuning during the early stages of multicellular evolution.

Despite their key housekeeping functions, chaperone expression is readily tunable. For instance, different animal tissues display distinctive chaperone expression profiles that specify their protein folding capacity (*47–49*). In yeast, Hsp90 chaperoning capacity surpasses demand under normal conditions as cells are viable with as little as 5% of wildtype Hsp90 under optimal growth conditions (*23*). Given the possibility for plasticity in Hsp90 expression, one possible alternative explanation for our results is that Hsp90 downregulation is not directly linked to multicellular adaptation; instead, it might be a by-product of metabolic changes or other mutations that occurred during the MuLTEE. This is unlikely however, given that Hsp90 downregulation was observed in all five replicate populations that evolved macroscopic size, restoring Hsp90 levels to the ancestral level reversed the cellular elongation phenotype and reduced multicellular fitness, and Hsp90 expression did not decrease in a parallel evolutionary treatment which remained microscopic. Instead, our data indicate that the downregulation of Hsp90 derives from decreased transcriptional activity of Hsf1. Although Hsf1 expression was unaltered, its activity can be modulated by post-translational modifications that affect its DNA binding without affecting its expression (*50–52*), which is consistent with what was observed in the T600 cells. Finally, while we provide evidence that the cellular elongation phenotype driven by reduced Hsp90 is mediated through its client Cdc28, we also note that Hsp90 has hundreds of other clients. It is therefore plausible that altered chaperoning of other proteins may contribute to the emergence of additional multicellular phenotypes.

Our results have several implications for the continued evolution of multicellularity in the MuLTEE. First, the evolution of increased cell aspect ratio via cell cycle delay may help entrench multicellularity. Entrenchment is a process through which multicellular traits become stabilized over evolutionary time, reducing the likelihood of reversion to unicellularity (*53*). We show here the reduction of Hsp90 promotes multicellular fitness by mediating cellular elongation via a delayed cell cycle. However, at the single-cell level, delaying progression through the cell cycle should be costly, as unicellular fitness in laboratory populations is primarily dependent on a genotype’s growth rate (*54*). Second, increased cellular aspect ratio was not driven exclusively by Hsp90– macroscopic lineages possessed an average of 32 mutations by T600, many of which are in genes affecting the cell cycle, filamentous growth, and budding processes (*19*). Mutations in some of these genes increase cellular aspect ratio and group size independently from Hsp90 (*19, 55*), and future work will be required to explore the joint evolution of genetic and epigenetic mechanisms underpinning the origin and maintenance of novel multicellular traits. Lastly, it will be interesting to further examine the permanence of the altered heat shock response, Hsp90 expression, and their consequences on protein evolution and stress tolerance over long evolutionary time scales (*56*).

Overall, our study shows how Hsp90, a key protein chaperone in all eukaryotes, influences the evolution of multicellularity in *S. cerevisiae*. By modulating the proteostasis of its client proteins, Hsp90 can generate novel multicellular phenotypes that are adaptive and heritable. This reveals how protein folding systems can shape emerging multicellular genotype-phenotype relationships, supporting the progression of an evolutionary transition in individuality. Our approach, using long-term open-ended experimental evolution, opens new avenues for fundamental biological discovery, and highlights the importance of epigenetic mechanisms in the transition to multicellularity.

## Materials and Methods

### Methods

#### Snowflake yeast strains and long-term evolution experiment

All yeast strains are derived from the Y55 diploid background and are listed in supplementary table 1. The ancestor strain was generated by deletion of both copies of the *ace2* transcription factor (*ace2Δ::KANMX/ace2Δ::KANMX*, or *ace2Δ*) and selecting a randomly generated petite mutant (i.e., carrying deletions in its mitochondrial DNA that render it non-functional). This anaerobic yeast snowflake strain was further subjected to a long term evolution experiment (*18*). In brief, five replicate populations of Ancestor snowflake were grown in 10 mL of YEPD media (1% yeast extract, 2% peptone, 2% dextrose) for 24 h, using settling selection to select for larger cluster size. Every 24 h over 600 consecutive days, 1.5 ml of culture were transferred into 1.5 mL Eppendorf tubes, and left on the bench for 3 minutes to allow cell settling. The bottom 50 µl of the settled culture was then transferred into fresh 10 mL of YPD for the next round of 24h growth and settling selection.

#### Experimental conditions and treatments

Prior to experiments, cells were grown in YPD medium (1 % yeast extract, 2% Bacto peptone, 2% glucose) for 24h and subject to daily settling selection. In total, three rounds of growth and settling selection was applied. On the fourth day, cells were subject to settling selection, grown for 4-6 h to exponential phase then experiments conducted. For heat shock experiments, cells were incubated at 42°C for 30 min, and then allowed to recover at 30°C for 10 min and RT for 5 min.

For treatment of Ancestor cells with radicicol, cells were incubated at 30°C with a final concentration of 40 µm of radicicol (solubilized in DMSO) for 6 h. Control cells were incubated with equivalent volume of DMSO.

#### Transformation of snowflake yeast

Ancestor, T200, and T400 snowflake yeast strains were transformed via a standard LiAc method (*57*). Cells were grown in exponential phase in YPD medium (1 % yeast extract, 2% Bacto peptone, 2% glucose) for 24 h. After 24 h, 250 µls of biomass (per transformation) was transferred to 10 mL of YPD and grown for 4 h. Cells were then pelleted via centrifugation at 1000 *g* for 1 min and washed once with H_2_O and once with 10 mM LiAC. Cells were pelleted and resuspended in 240 μl of PEG buffer (40% PEG-3350 (m/V), and mixed with 36 μl of 1M LiAC, plasmid DNA (1 µg) or PCR product (10μl) and 10 μl of 100 mg/ml salmon sperm DNA (prior to use boiled for 10 min at 100 °C and cooled down on ice). Yeast were then subjected to heat shock at 42°C for 30 min and spun down at 300 *g* for 5 min. Cells were incubated in 3 ml of YPD media for 3 h and then plated on YPD plates with appropriate antibiotics. Snowflake yeast strain T600 were transformed via electroporation. Cells were grown in YPD medium (1 % yeast extract, 2% Bacto peptone, 2% glucose) for 24 h. After 24 h, 100 µls of biomass (per transformation) were transferred to 10 mL of YPD and grown for 4 h. Cells were then pelleted by centrifugation at 1000 *g* for 5 min, washed twice with H_2_O and once with 1M sorbitol. Cells were incubated in 10 ml of conditioning solution (0.1M LiAC/10 mM DTT) at 30°C for 30 min. A cell pellet was collected by centrifugation at 1000 *g* for 5 min, then washed twice with 1M sorbitol. Cells were resuspended in 350μl of electroporation buffer (1M Sorbitol/1mM CaCl_2_) and mixed with plasmid DNA (1µg) or PCR product (10μl). The transformation mixture was transferred to electroporation cuvettes (Biorad), and electroporation conducted at 2.5kV (Biorad micropulser). Cells were transferred to 5ml of YPD to recover overnight, then plated on plated on YPD plates with appropriate antibiotics.

#### Protein extraction and Western Blot

Cells were harvested by centrifugation at 1000 *g* for 5 min and the pellet washed once with H_2_O and once with PBS. Cells were resuspended in 800 μl of PBS and 200 μl of 50 % Trichloroacetic acid (TCA), and stored for 30 min at −80°C to allow protein precipitation. TCA treated cells were pelleted, washed twice with ice-cold 80% acetone and air-dried. Pellets were dissolved in 180 μl of lysis buffer (1% SDS, 0.1M sodium hydroxide) and boiled for 5 min. Protease inhibitors were then added (Roche). Quantification of protein concentration was done using a BCA Protein assay Kit (Thermo Fisher Scientific). For western blot, 5-50 µg of protein was mixed with 6x loading buffer (48% glycerol and 0.03% bromophenol blue) and run through SDS-PAGE gel. For detection, primary mouse Anti-Hsp90 (StressMarq) was used 1:1000, primary mouse Anti-Cdc28 (Santa Cruz Biotechnlogy, inc) was used 1:1000 and for loading control, mouse Anti-β-actin (Abcam) was used 1:1000. For secondary antibody, horseradish peroxidase (HRP) conjugated Rabbit Anti-Mouse (Invitrogen) was used 1:2000.

#### RNA extraction and RT-qPCR

Cells were harvested by centrifugation at 1000 *g* for 5 min. Cell pellets were resuspended in 800 μl of TRI reagent (Zymo research) then transferred to 2 ml Touch Micro-Organism Lysing Mix tubes (Omni). Cells were then mechanically disrupted with Precellys 24 Tissue Homogenizer (Bertin Instruments) at 6000 rpm for 15s for a total of 4 cycles with a 2 min break on ice between cycles. RNA was purified using a Direct-zol RNA Miniprep Kit (Zymo Research) and following the manufacturer’s instructions. RNA concentration and purity was measured by NanoDrop, and RNA samples were stored at −80°C. cDNA was prepared from purified RNA (1 µg) using Oligo(dT) (Invitrogen) and reverse transcriptase Superscript IV (Invitrogen) according to the manufacturer’s instructions. Synthesized cDNA was diluted 1:10 and 5 µl mixed with SsoAdvanced Universal Sybr Green supermix (Biorad). qPCR was performed using the Biorad CFX96 PCR machine. Relative RNA levels were quantified using the ΔΔCT method. *ACT1* was used as the normalization control.

#### RNA sequencing and data analysis

RNA for sequencing was extracted as described above. RNA-seq library preparation and sequencing was performed by the Sequencing Unit at the Institute of Molecular Medicine Finland (FIMM) Technology Centre, University of Helsinki. Illumina TruSeq Stranded mRNA library preparation Kit (Illumina) and NextSeq 500 Mid Output Kit PE75 (120 M reads) (Illumina) were used following manufacturer’s instructions. Quality control of raw reads were performed with FastQC v0.11.9, followed by read filtering with Trim Galore v0.6.6 and rRNA removal with SortMeRNA v4.2.0. Transcriptome mapping was done with Salmon v1.4.0 and quality control of read mapping was done with STAR v2.7.8a and Qualimap v2.2.2d. Summary of quality control results was reported with MultiQC v1.9. Differentially expressed genes between Ancestor and T600 cells were found using DESeq2 v1.32.0 (adjusted p-value < 0.05, Wald test) in R 4.1.0. Functional enrichment of differentially expressed genes by over representation analysis was done with clusterProfiler v4.0.0. For this study, the yeast genome/transcriptome references and gene annotations from Ensembl release 103 were used, which are based on yeast S288C genome assembly R64-1-1.

#### Hsf1 luciferase activity assay

Snowflake yeast strains were transformed with genomically-integratable linearized plasmid, pCYB2194-3xHSE-NLucPEST-HygMX, which is based on the plasmid pAM17 (HYG – P_cyc1-3xHSE-_yNlucPEST) (a generous gift from Claes Andréasson / Uni. Stockholm). Cells were grown in standard experimental conditions and induction of Hsf1 was done with heat shock at 42°C for 10 min. Cells were washed once with H_2_O and then mixed with 400 µl of lysis buffer (50 mM TrisHCl ph7.5, 300 mM NaCl and 10 mM Imidazole). Proteins were extracted by mechanical disruption with Precellys 24 Tissue Homogenizer (Bertin Instruments) at 6000 rpm for 15s for a total of 4 cycles with 2 min break on ice between cycles. Protein concentration was measured using a Nanodrop (Thermo Fisher Scientific) and a BCA protein assay kit (Thermo Fisher Scientific). Nano-Glo substrate (Promega GmbH, Germany) was prepared at 1:100 substrate to buffer, and mixed 1:10 (20 µl) with lysate (180 µl at a of concentration 0.5 µg/ml) in a 96-well plate. Bioluminescence was measured immediately, using an Enspire Plate Reader (Perkin Elmer).

#### Hsf1 ChIP-QPCR assay

Ancestor and T600 cells expressing endogenously tagged Hsf1-GFP were grown in standard experimental conditions. Cells were then treated with formaldehyde (1% final concentration) for 15 min at RT for protein crosslinking. The crosslinking reaction was quenched by adding glycine (final concentration 125 mM) and incubating for 5 min. Cells were collected by centrifugation at a 1000 *g* for 5 min at 4°C. The supernatant was removed, pellet re-suspended in 10 ml of TBS pH 7.5, and then centrifuged again at 1000 *g* for 5 min at 4°C. Cells were washed with ice cold H_2_O, pelleted and resuspended in 800 µl of lysis buffer (50 ml 1 M Hepes/KOH (pH 7.5), 28 ml 5 M NaCI, 2 ml 500 mM EDTA, 100 ml 10% Triton X-100, 1 g Na-deoxycholate, Protease inhibitor cocktail tablet (Roche) and 1 mM PMSF). Cells were lysed with the Precellys 24 Tissue Homogenizer (Bertin Instruments) at 6000 rpm for 15s for a total of 4 cycles with a 2 min break on ice between cycles. Lysates were recovered and subjected to sonication using a Bandolin Sonopuls sonicator (10 cycles with 30s pulse). Cell debris was removed by centrifugation at 3000 rpm for 5 min at 4°C. To perform immunoprecipitation, the equivalent of 500-800 µg of protein was incubated with 1 µg of anti-GFP antibody (Sigma-Aldrich) and then incubated with Protein A/G magnetic beads (Thermofisher) overnight at 4°C. Beads were washed with lysis buffer and eluted with TE/1%SDS (pH 8.0). Crosslinks were reversed by incubation at 65°C, and elutants were treated with RNAse A and proteinase K. DNA was purified using a GeneJET PCR purification kit (Invitrogen). Immunoprecipitated DNA was mixed with SsoAdvanced Universal Sybr Green supermix (Biorad) and qPCR conducted using the Biorad CFX96 PCR machine. Primers against the HSE regions of *HSC82* and *HSP82* were used. DNA levels were quantified as a percentage of input. Background was determined by signal arising from incubating an equivalent volume of chromatin extract without antibody. Signal from input was used to normalize against variation in yield of chromatin.

#### Microscopy

For still images, cells were stained with 100 μg/ml calcofluor for 5 min. Cells were washed twice with H_2_O then resuspended into 1 ml of H_2_O. 10 µl of cell suspension was dropped onto a slide and coverslip was firmly placed over the top. For time lapse experiments, cells were grown in YPD for 24 h then transferred to synthetic complete (SC) media and grown for 2 h. Clusters were then broken via mechanical disruption between two glass slides, and transferred to 96 well thin glass bottom imaging plate pre-coated with concanavalin A (Sigma Aldrich) in a volume of 200 μl of SC media. Well coating was performed by adding 100 μl of concanavalin A (2 mg/ml in PBS), incubating in room temperature for 30 min and washing wells twice with PBS. Imaging was performed with a customized Olympus IX-73 inverted widefield fluorescence light microscope DeltaVision Ultra (GE Healthcare) equipped with Pco edge 4.2ge sCMOS camera and CentOS 7 Linux operating system. Images were taken at 60x oil objectives and, depending on the fluorophore properties, Blue (ex. 390 nm – em. 435 nm) and Green (ex. 475 nm – em. 525 nm) filter settings. Imaging data was analyzed with FIJI ImageJ software.

#### Competition Assay

We measured the fitness of Hsc82 overexpression strains over three transfers of growth and settling selection. To hedge against possible off-target effects of genomic transformation, we replicated this experiment four times, with four independently-generated Hsc82OE clones. In each comparison, we competed four replicate populations of T600-GFP against an unmarked Hsc82OE strain. Prior to the competition, we inoculated each strain into 10 mL of YPD, and grew them for five days of growth and settling selection to allow each population to reach cluster size equilibrium. Each strain was then mixed in equal volumes with T600-GFP, and 50 μL of this mixture was inoculated into 10 mL of YPD media to start the competition. Every 24 h for 3 days, 1 mL of each competing population was transferred into 1 mL centrifuge tubes to perform settling selection. After 3 minutes of settling on the bench, the top 950 μL was discarded. The remaining 50 μL was transferred into 10 mL of fresh YPD media. The cellular biomass of GFP-tagged and non-GFP-tagged strains were estimated based on the cluster area with the following equation:

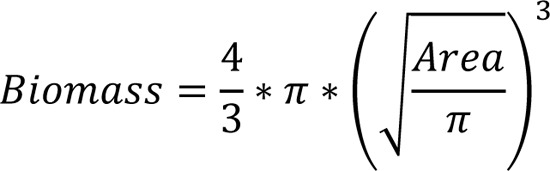

We used the biomass of GFP-tagged and non-GFP-tagged strains of initial mixtures and 3-day cultures to calculate fitness as a selection rate constant, using the method described by Richard Lenski here: https://lenski.mmg.msu.edu/ecoli/srvsrf.html. To account for the costs of GFP expression, we normalized the selection rate constant of T600-Hsc82OE clones by the selection rate constant of T600 vs T600-GFP (all fitness values shown in Figure 2F are normalized in this way).

#### Statistical Analysis

The statistical analyses tests used in this study were the two-tailed Student’s *t* test, one-way analysis of variance (ANOVA), two-way analysis of variance (ANOVA) and a nested analysis of variance (ANOVA). The analysis was conducted using the Prism statistical software program (version 9.0; GraphPad Software, Boston, MA, USA). A *p* value less than 0.05 was considered statistically significant. In the figure legends, *, **, ***, and **** indicate *P* values less than 0.05, 0.01, 0.001, and 0.0001, respectively. All experiments were performed with three replicates unless specified, and the error bars in the figure legends represent means ± SEM.

## Materials

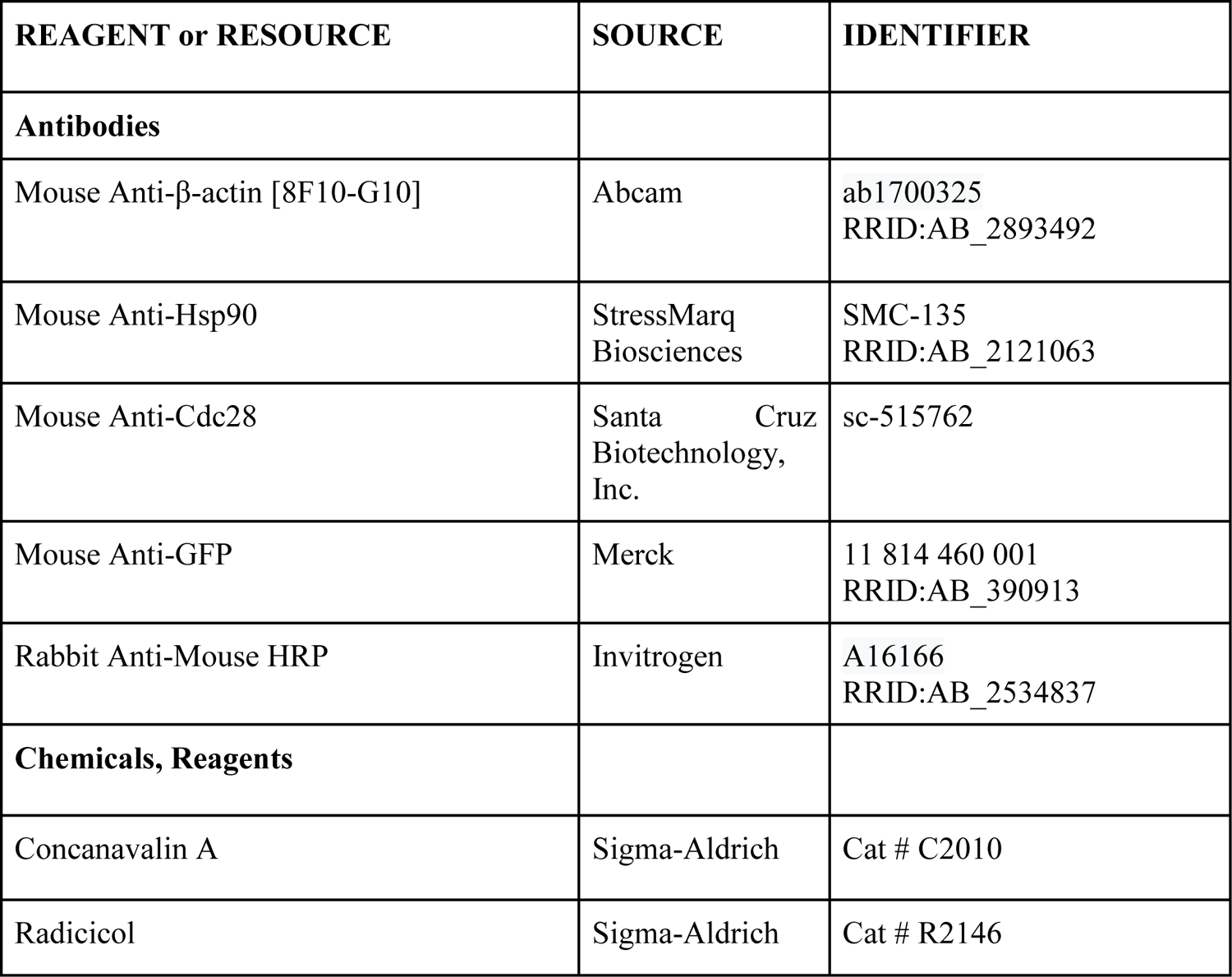
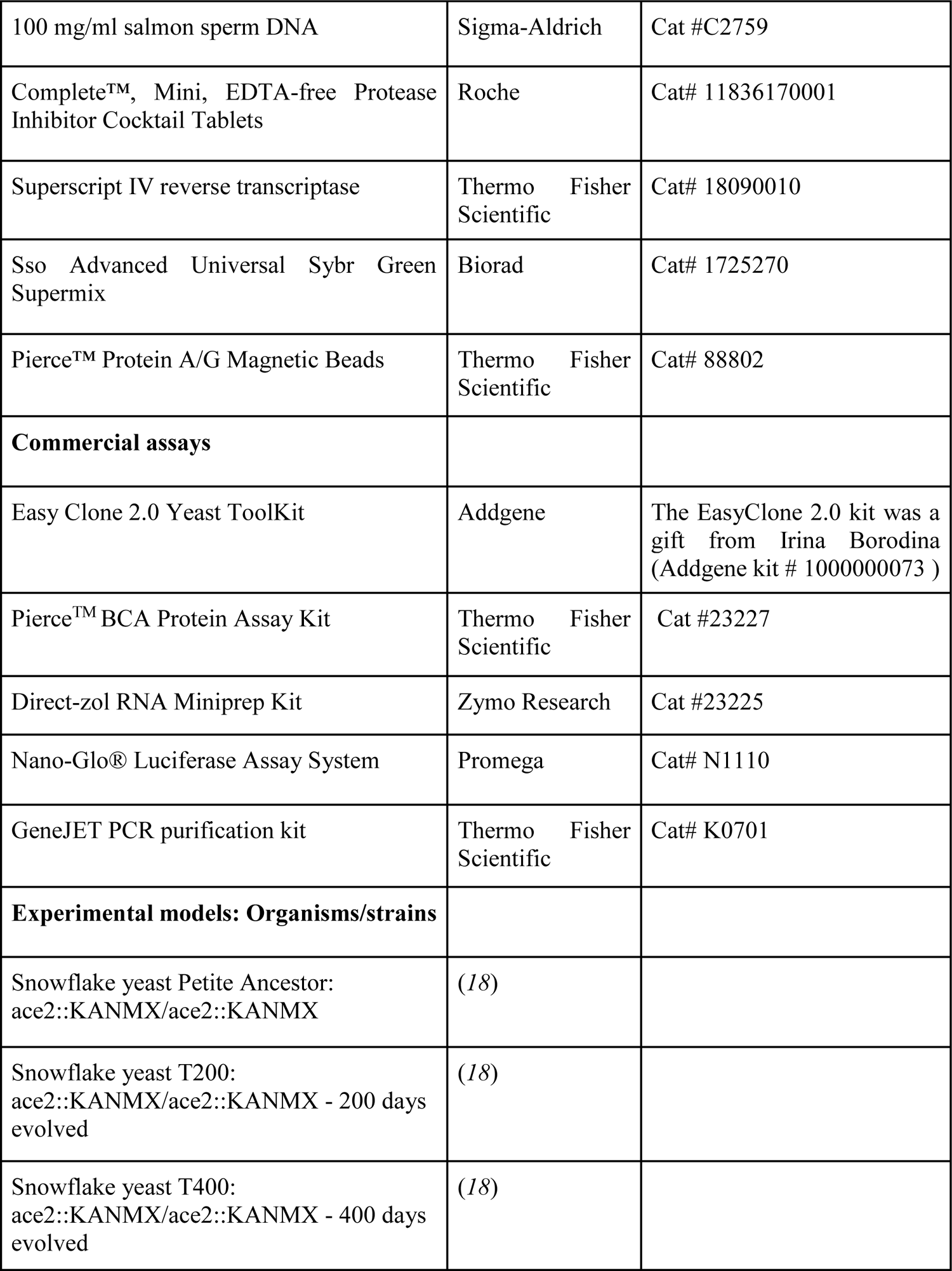
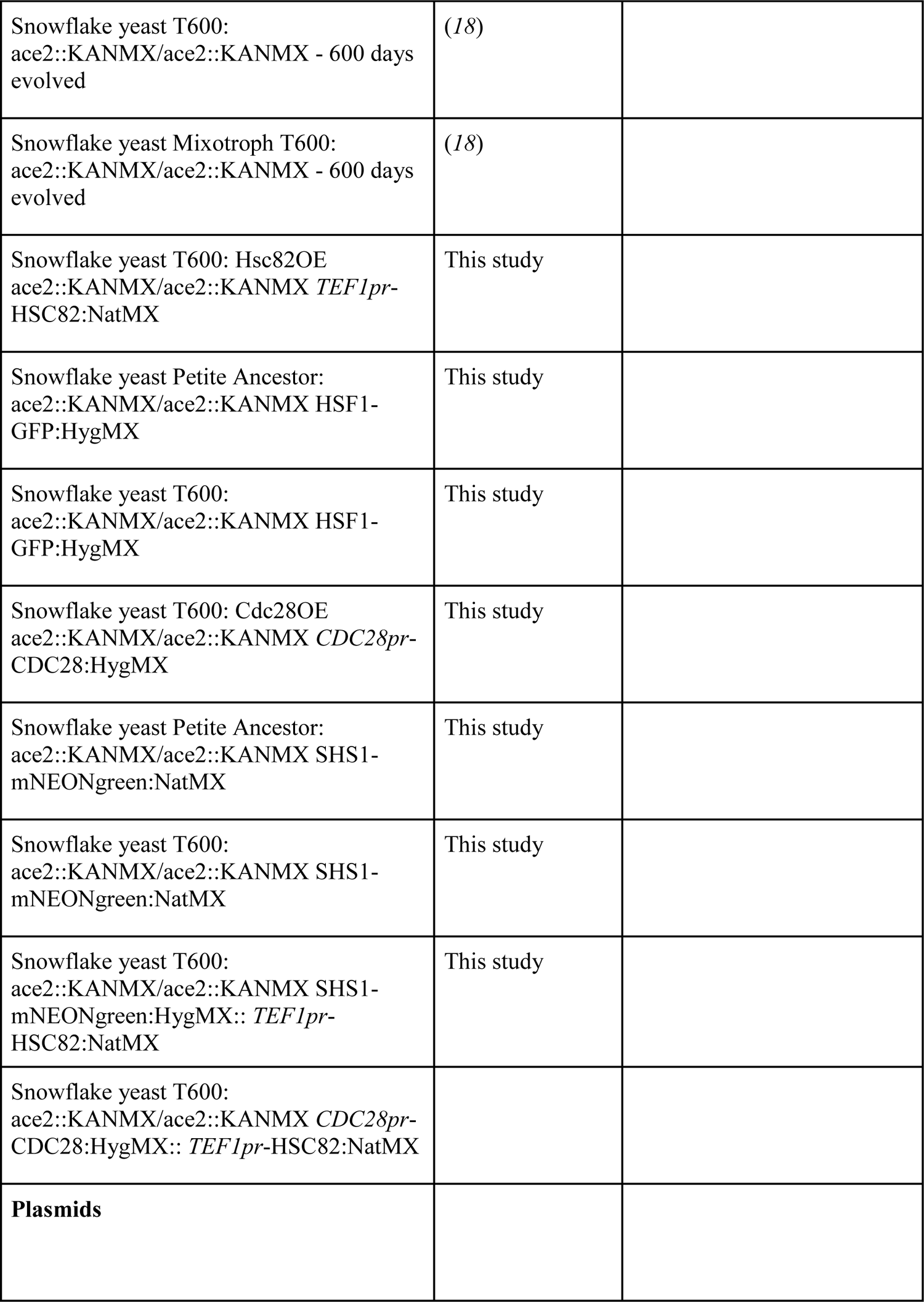
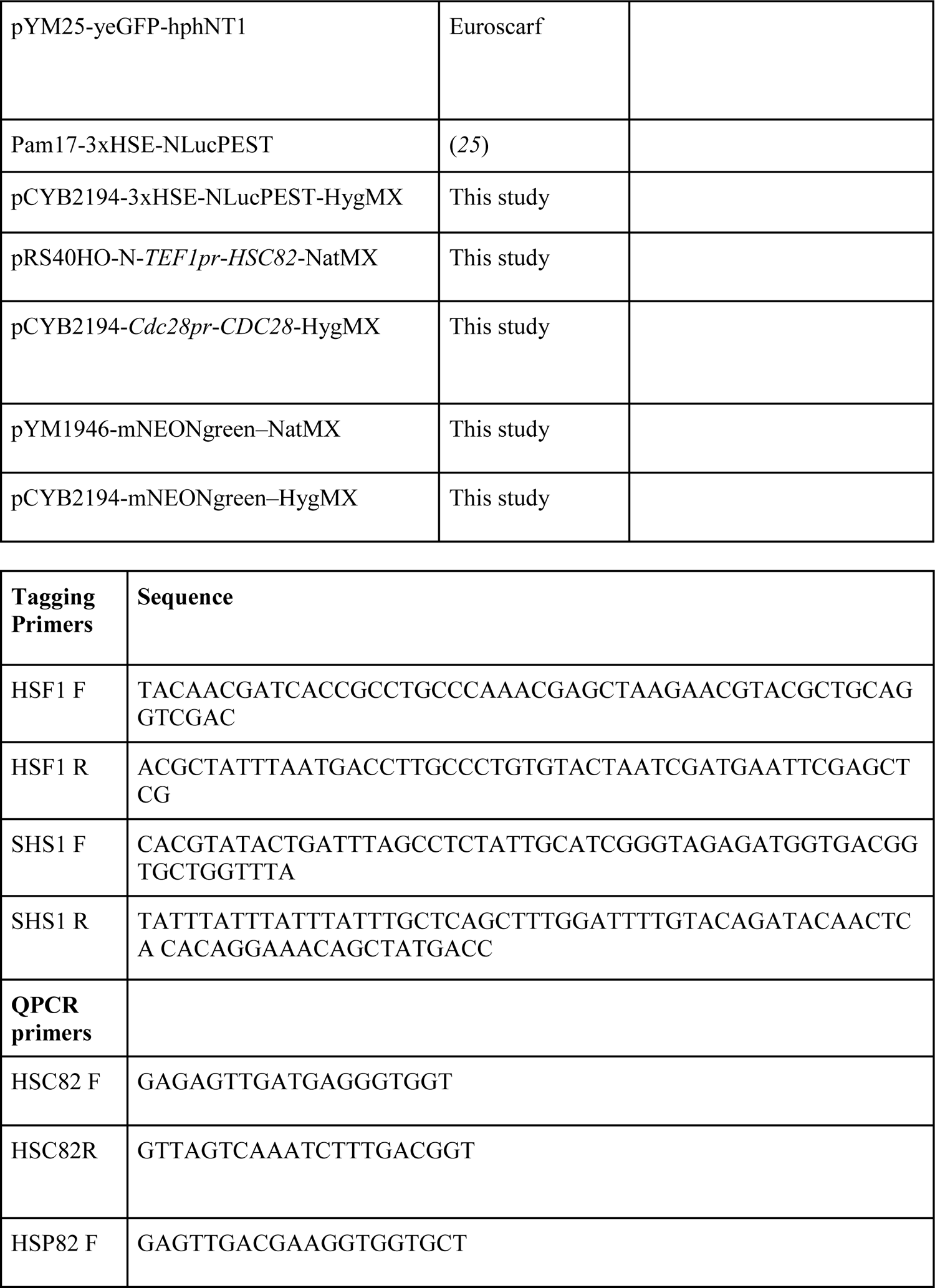
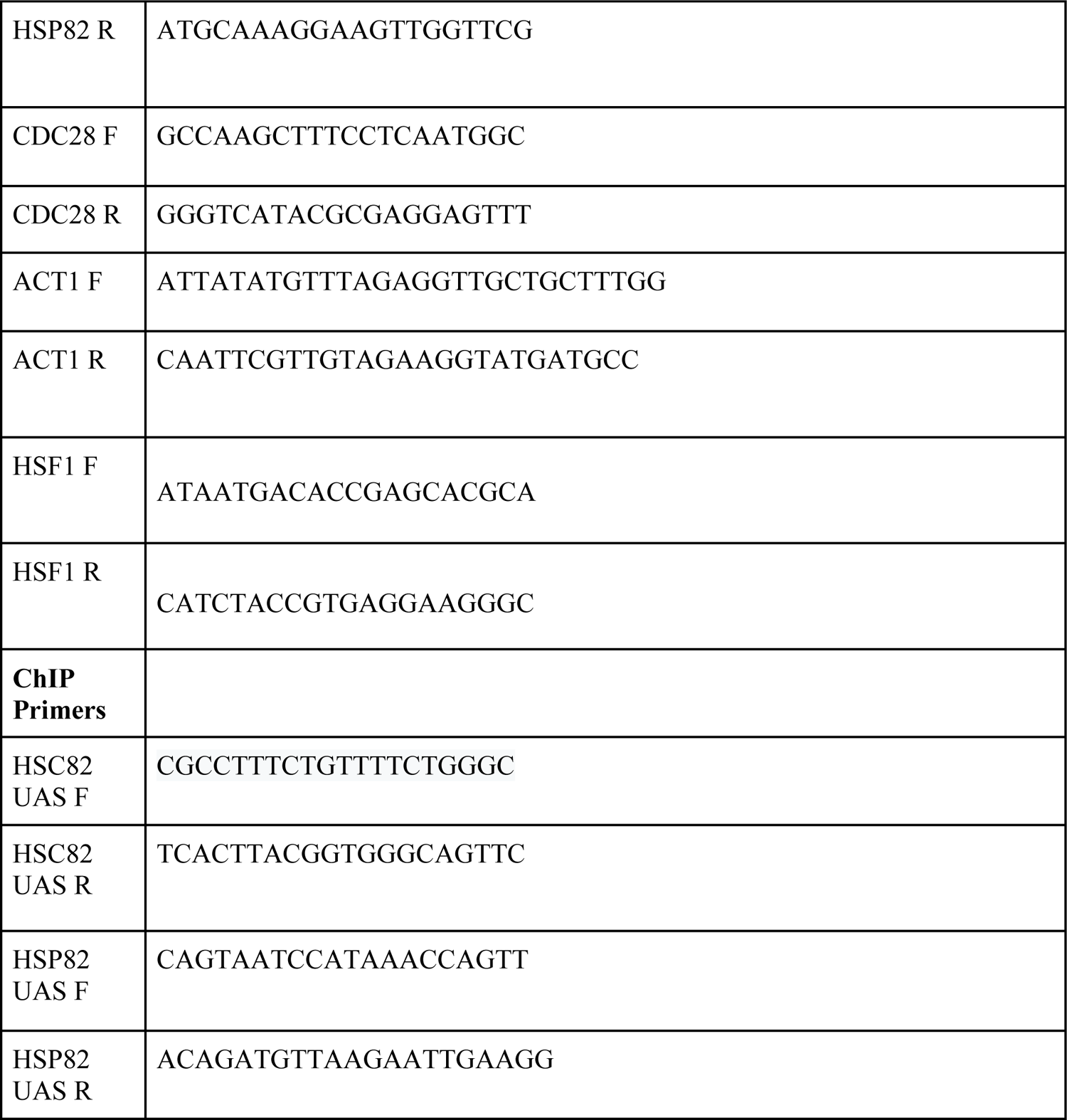

## Acknowledgments

We thank the members of the Saarikangas lab and Ratcliff lab for input and Claes Andréasson and David Gross for reagents and advice. We are grateful to the assisting personnel at the Light Microscopy (LMU), FIMM Genomics and DNA sequencing and Genomics (BIDGEN) units, as well as MIBS and BI Media Kitchens at the University of Helsinki. Graphics in Fig. 1D, 2E, 4F and 4H, were made in ©BioRender - biorender.com.

## Author contributions

Conceptualization: KM, WCR, JS

Methodology: KM, DTL, AJB, KT, GOB, MH, WCR, JS

Investigation: KM, DTL, AJB, KT, GOB, MH

Funding acquisition: WCR, JS

Supervision: WCR, JS

Writing and editing: KM, DTL, AJB, KT, GOB, MH, WCR, JS

## Data and materials availability

All data are available in the main text or the supplementary materials.

## Supplementary Materials

Figs. S1 to S4

References (18, 25, 57)

