## Supplemental Figures for "Proteostatic tuning underpins the evolution of novel multicellular traits"

Kristopher Montrose *et al*

**This PDF file includes:**

Figs. S1 to S4
References (*18, 25, 57*)

S1

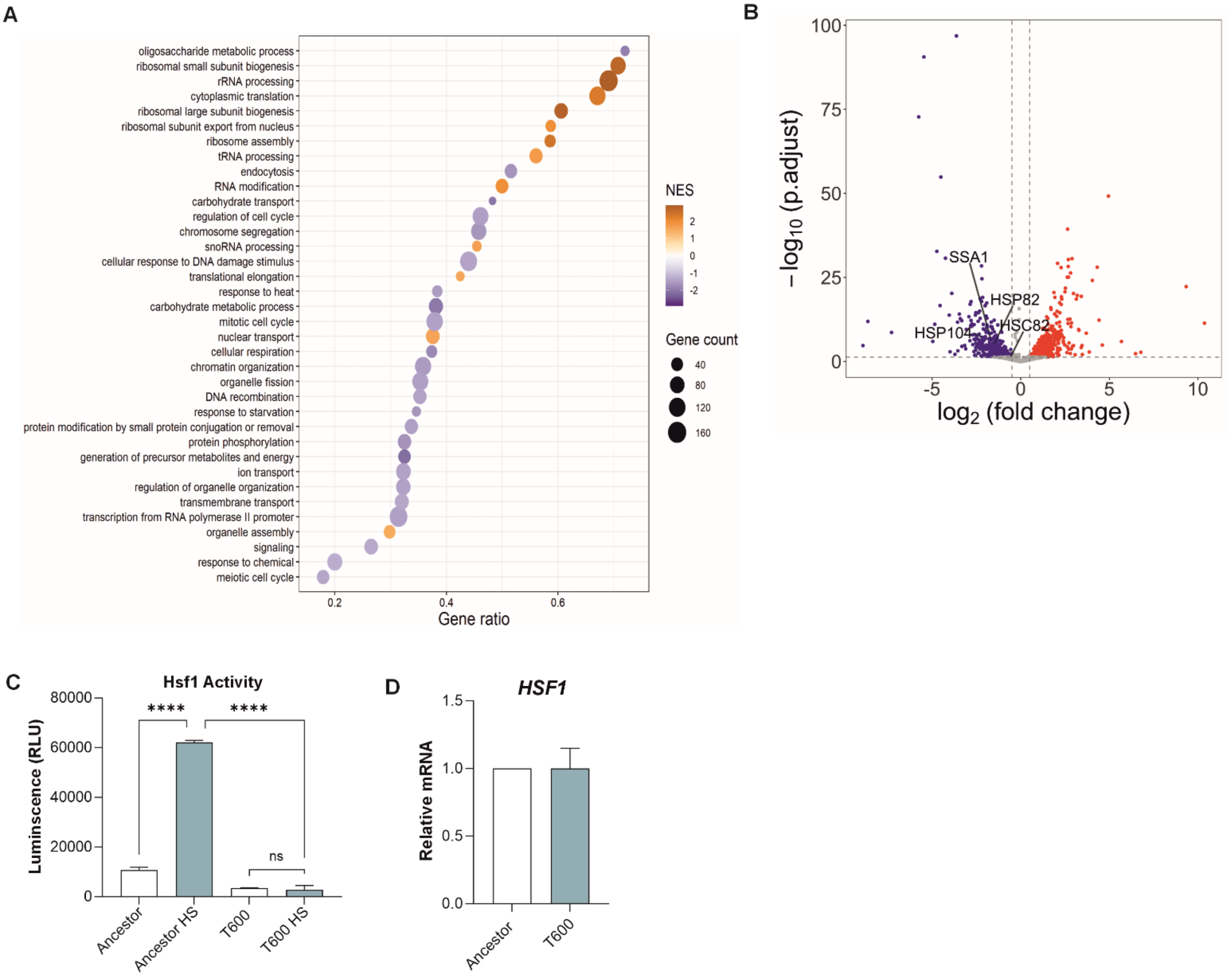


**Supplementary 1**

(A) Dot plot of differential gene expression between Ancestor and T600 snowflake yeast organized into the top 50 gene ontology hits ($n$=3). (B) Volcano plot of differential gene expression between Ancestor and T600 highlighting changes in chaperone proteins Hsp90 (Hsc82 and Hsp82), Hsp70 (Ssa1) and Hsp104 ($n$=3). (C) Hsf1 activity before and after heat shock in Ancestor and T600 cells measured as luminescence. Luminescent readings of substrate alone were subtracted as background ($n$=4, $F_{3, 12}$ = 686.1, $p$ < 0.0001, one way ANOVA, *Tukey's* post hoc test Ancestor vs Ancestor HS $p$ < 0.0001, T600 vs T600 HS $p$ = 0.97, Ancestor HS vs T600 HS $p$ < 0.0001). (D) Quantification of *HSF1* expression level by RT-qPCR of T600 compared to Ancestor ($n$=6, $t$ = 0.0071, $p$ = 0.994, two-sample t-test). All values represent mean ± SEM.

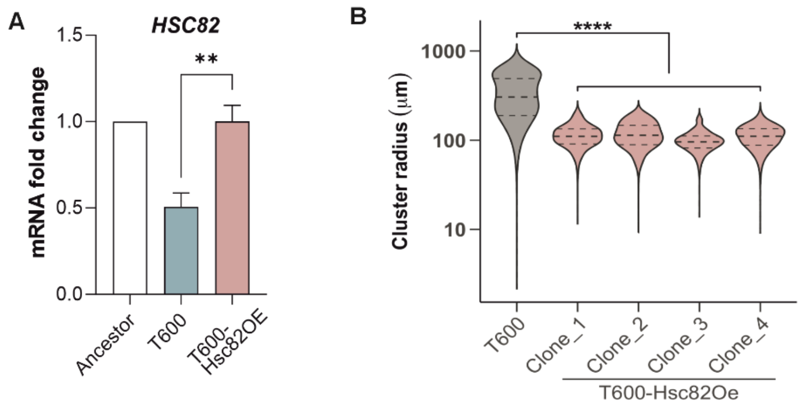


**Supplementary 2**

(A) Quantification of *HSC82* expression level by RT-qPCR in T600 and T600-Hsc82OE compared to Ancestor ($n$=4, $F_{2,6} = 15.96$, $p = 0.004$, one way ANOVA, *Tukey's* post hoc test T600 vs T600-Hsc82OE $p = 0.006$). (B) Cluster size as a measure of cluster radius (µm) for T600 and four clones of T600-Hsc82OE ($F_{1,6226} = 3024$, $p < 0.0001$, one way ANOVA, *Tukey's* post hoc test T600 vs T600-Hsc82OE 1-4 $p < 0.0001$). All values represent mean ± SEM.

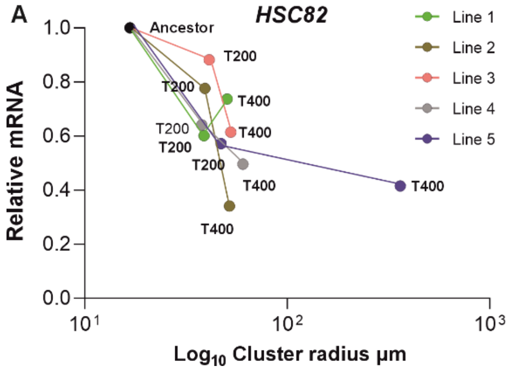


**Supplementary 3**
(A) Scatter plot of *HSC82* expression against cluster radius for T200 and T400 cells for each of the five lines of aerobic snowflake yeast. ($r$ = 0.66, $p$ = 0.02, $y$ = -0.40$x$+1.33, Linear regression).

S4

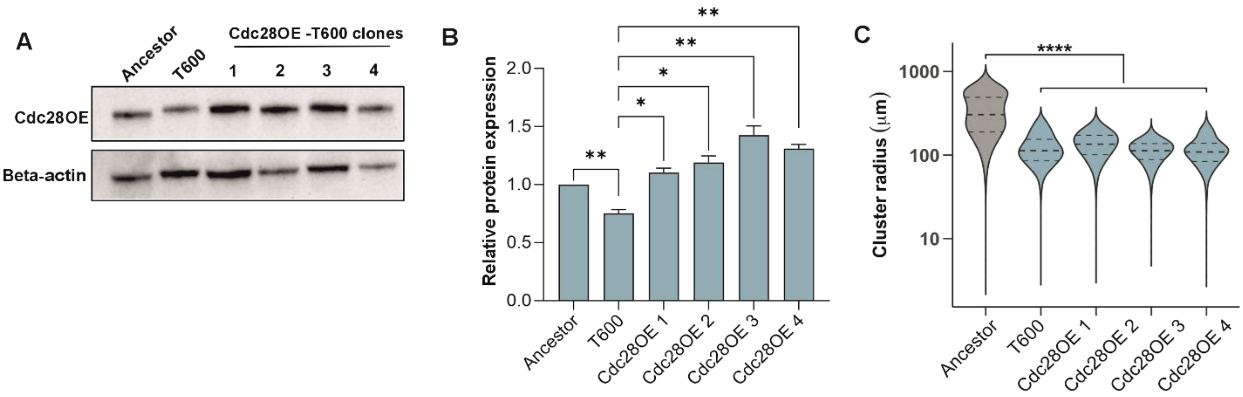


**Supplementary 4**

(A) Representative immunoblot of Cdc28 expressed by Ancestor, T600 and 4 clones of T600-Cdc28OE, detected with anti-Cdc28 antibody. Antibody against Beta-actin was used as a loading control. (B) Quantification of the band intensity of Cdc28 in T600 and four clones of T600-Cdc28OE relative to the Ancestor. Bands were normalized to loading control (*n*=5, $F_{2,10} = 27.37$, $p < 0.0001$, one way ANOVA, *Tukey's* post hoc test Ancestor vs T600 $p = 0.0081$, T600 vs T600-Cdc28OE 1 $p = 0.0251$, T600 vs T600-Cdc28OE 2 $p = 0.0116$, T600 vs T600-Cdc28OE 3 $p = 0.006$, T600 vs T600-Cdc28OE 4 $p = 0.002$). (C) Cluster size as a measure of cluster radius (μM) for T600 and four clones of T600-Cdc28OE ($F_{1,7195} = 3477$, $p < 0.0001$, one way ANOVA, *Tukey's* post hoc test T600 vs T600-Cdc28OE 1-4 $p < 0.0001$).
